# RNA Pol IV has antagonistic parent-of-origin effects on Arabidopsis endosperm

**DOI:** 10.1101/2021.03.24.436774

**Authors:** Prasad R.V. Satyaki, Mary Gehring

## Abstract

Gene expression in endosperm – a seed tissue that mediates transfer of maternal resources to offspring – is under complex epigenetic control. We show here that plant-specific RNA Polymerase IV mediates parental control of endosperm gene expression. Pol IV is required for the production of small interfering RNAs that typically direct DNA methylation. We compared small RNAs, DNA methylation, and mRNAs in *A. thaliana* endosperm from reciprocal heterozygotes produced by crossing wild-type plants to Pol IV mutants. We find that maternally and paternally acting Pol IV have divergent effects on endosperm. Losses of maternal and paternal Pol IV impact sRNAs and DNA methylation at distinct genomic sites. Strikingly, maternally and paternally-acting Pol IV have antagonistic impacts on gene expression at some loci, divergently promoting or repressing endosperm gene expression. Antagonistic parent-of-origin effects have only rarely been described and are consistent with a gene regulatory system evolving under parental conflict.

## Introduction

Parents influence zygotic development in viviparous plant and animal species. In flowering plants, parent-of-origin effects on offspring development are observed in an embryosurrounding seed tissue called the endosperm (Gehring & Satyaki, 2017). Endosperm does not contribute genetic material to the next generation but mediates maternal nutrient transfer to the embryo, coordinates growth between the embryo and maternal tissues, sets seed dormancy and regulates germination, and acts as a nutrient store to support seedling growth (Jing Li & Berger, 2012). Endosperm is typically triploid and develops from the fertilization of a diploid female gamete, called the central cell, by one of two haploid sperm cells that are released by pollen. Violations of the balanced ratio of two maternal to one paternal genomes disrupts normal endosperm development in a parent-of-origin dependent manner (Milbocker & Sink, 1969; Müntzing, 1936; Povilus et al., 2018; Scott et al., 1998; Stoute et al., 2012). In some *A. thaliana* accessions, crosses between tetraploid mothers and diploid fathers exhibit reduced endosperm proliferation and smaller mature seeds while reciprocal crosses where the fathers are tetraploid (paternal excess crosses) exhibit prolonged endosperm proliferation and larger or aborted seeds. These parent-of-origin effects on endosperm development have been interpreted under the aegis of the parental conflict or kinship model (Haig, 2013). According to this model, when a mother mates with more than one father, the inclusive fitness of the mother is optimal when her resources are equally distributed among her progeny. The inclusive fitness of the father is optimal when his progeny are able to acquire more maternal resources than other half-siblings. Such conflicts are postulated to lead to arms races whose impacts may be observed in the molecular machinery mediating parental control. However, our understanding of the impact of conflict on endosperm biology is limited by our incomplete understanding of molecular and genetic mechanisms guiding parental control of endosperm development.

Recent data indicate that mutations in RNA Polymerase IV have effects on reproduction, endosperm, and seed development in multiple species (Erdmann et al., 2017; Grover et al., 2018; Kirkbride et al., 2019; Martinez et al., 2018; Satyaki & Gehring, 2019; Wang et al., 2020). RNA Pol IV functions as part of the RNA directed DNA methylation (RdDM) pathway, in which it produces relatively short, non-coding transcripts that are converted into double stranded RNA by RDR2 (Blevins et al., 2015; S. Li et al., 2015; Zhai et al., 2015). These double-stranded RNAs are cleaved into 24nt small RNAs (sRNAs) by DCL3 and single strands are loaded into ARGONAUTE proteins that help target the *de novo* DNA methyltransferase DRM2, which acts in conjunction with RNA Pol V and several other proteins, to methylate DNA (Matzke & Mosher, 2014). *NRPD1*, which encodes the largest subunit of RNA Pol IV, has roles in endosperm gene dosage control. Endosperm gene expression typically reflects the ratio of two maternally and one paternally inherited genomes, such that for the majority of genes approximately two-thirds of genic transcripts are derived from maternal alleles (Gehring et al., 2011; Pignatta et al., 2014). A survey of allele-specific gene expression in *nrpd1* mutant endosperm found that Pol IV is required to maintain the 2:1 maternal to paternal transcript ratio in the endosperm and that loss of Pol IV leads to the mis-regulation of several hundred genes (Erdmann et al., 2017). Additionally, loss of function mutations in *NRPD1* or other members of the RdDM pathway can repress seed abortion in crosses of diploid mothers and tetraploid fathers (Erdmann et al., 2017; Martinez et al., 2018; Satyaki & Gehring, 2019). In *B. rapa*, loss of *NRPD1*, *RDR2*, or *NRPE1* results in high rates of seed abortion due to maternal sporophytic effects (Grover et al., 2018). Loss of Pol IV in both *B. rapa* and in *A. thaliana* also results in smaller seed sizes (Grover et al., and RNA Pol IV is essential to post-meiotic pollen development in *C. rubella* (Wang et al., 2020).

Molecular data point to the intriguing possibility that mutations in RNA Pol IV have parent-of-origin effects on endosperm. A comparison of sRNAs in wild-type whole seeds (which includes maternal seed coat, endosperm, and embryo) with *NRPD1*+/- endosperm from crosses where the mutation in *NRPD1* was either maternally- or paternally-inherited suggested that loss of maternal *NRPD1* affected more sRNA loci than the loss of paternal *NRPD1* (Kirkbride et al., 2019). Although the comparison of sRNAs from wild-type whole seeds to mutant endosperm in this study makes definitive conclusions difficult to draw, it raises the potential question of if and how the loss of *NRPD1* has parent-of-origin effects on sRNA production.

To examine the impacts of parental Pol IV activity on endosperm in more detail, we examined sRNA and mRNA transcriptomes in wild-type endosperm, *nrpd1* homozygous mutant endosperm, and *nrpd1* heterozygous endosperm where the mutant allele was inherited from a homozygous mutant mother or father. We also examined methylomes in wild-type and endosperm from the reciprocal heterozygotes. Analysis of these data demonstrate that maternal and paternal *NRPD1* have distinct parental effects on endosperm, some of which are antagonistic.

## Results

### Maternal Pol IV inhibits, whereas paternal Pol IV promotes, interploidy seed abortion

We tested if the molecular data supporting distinct functions for Pol IV in the mother and the father (Kirkbride et al., 2019) could be supported by genetic analyses. We reanalyzed previously published data (Erdmann et al., 2017) to specifically test the effects of the loss of maternal Pol IV *vs*. paternal Pol IV in the context of interploidy, paternal excess crosses (diploid mother pollinated by tetraploid father). Inheritance of a mutant *nrpd1* allele from diploid mothers resulted in 4% normal seed in a cross to tetraploid fathers, which was significantly different than 7.1% normal seed observed when wild-type diploid mothers are crossed to wild-type tetraploid fathers. Crosses between wild-type diploid mothers and tetraploid *nrpd1* fathers resulted in 64.8% normal seed. Paternal rescue by *nrpd1* was diminished when the diploid mother was also mutant, resulting in 37.5% normal seeds. Thus, we conclude that maternal *NRPD1* promotes interploidy seed viability and paternal *NRPD1* represses seed viability (Fig. S1). This is consistent with observations that paternal excess seed viability was promoted by the maternal activity of *DCL3* and repressed by the paternal activity of *DCL3* (*DCL3* functions downstream of *NRPD1*) (Satyaki & Gehring, 2019). Interploidy crosses are a sensitive genetic assay to detect endosperm phenotypic effects. However, paternal excess endosperm displays wide-spread transcriptomic changes (Satyaki & Gehring, 2019), which make it a poor system to understand the specific role of RNA Pol IV in endosperm development. Therefore, for all subsequent experiments we examined endosperm molecular phenotypes in the context of balanced crosses (diploid x diploid) where either one or both parents were homozygous mutant for the *nrpd1a-4* allele.

### Loss of maternal or paternal Pol IV activity impacts small RNAs at distinct sites

To determine the role of Pol IV in sRNA production in the endosperm, we first identified Pol IV-dependent endosperm sRNAs. Previously we showed that 24 nt sRNAs were the predominant sRNA species in endosperm and exhibited a broader distribution over genes and transposable elements (TEs) than in other tissues (Erdmann et al., 2017). We profiled small RNA populations in three replicates of endosperm derived from crosses of L*er nrpd1* females pollinated by Col-0 *nrpd1* males (7 days after pollination) and compared them with our previously published sRNA libraries from L*er* x Col-0 wild-type F1 endosperm (female parent in cross written first) (Erdmann et al., 2017) (Supplementary Table 1).

We identified 21,131 sRNA peaks in wild type endosperm using ShortStack (Axtell, 2013). 76.9% of these were predominantly populated by 24nt sRNAs, with 1.1%, 0.2%, and 2.2% of peaks dominated by 23, 22, or 21 nt sRNAs, respectively. An additional 19.7% of peaks were either dominated by a non-canonical sRNA size or had no predominant size class (Fig. S2A). The majority of sRNAs were genetically dependent on *NRPD1*, with 99% of 24nt sRNA peaks, 94.87% of 22nt sRNA peaks and 70.1% of 21nt sRNA peaks absent in *nrpd1*-/- endosperm (Fig. S2B). To enable downstream comparisons to expression, we binned sRNAs by size (21 to 24 nt) and calculated read counts overlapping TEs and genes encoding proteins, miRNA, and other ncRNA. We used DESeq2 (Love et al., 2014) to separately identify genes and TEs with significant differences in Pol IV-dependent sRNA populations. Consistent with the peak-based analysis, loss of RNA Pol IV abolished 21-24 nt small RNAs at most TEs and genes, while most miRNAs were not impacted (Supplementary Table 2). 21-23 nt sRNAs were often lost at the same loci as 24 nt sRNAs (Fig. S2C), suggesting that sRNAs of differing sizes arose from the same Pol IV transcript in the wild-type but were likely processed into RNAs shorter than 24 nt by different downstream DICERs or by the exosome components *Atrimmer1* and *2* (Daxinger et al., 2009; Ye et al., 2016). Pol IV-dependent 21-23 nt sRNAs have been identified in other tissues, indicating this finding is not specific to endosperm (Panda et al., 2020; Wang et al., 2020; Wu & Zheng, 2019).

After identifying Pol IV-dependent sRNAs, we asked whether loss of one parent’s Pol IV influenced the abundance of Pol IV-dependent sRNAs in *nrpd1* heterozygous endosperm. We sequenced small RNAs from two replicates of L*er* female x Col-0 *nrpd1*-/- male (referred to as pat *nrpd1*+/-) endosperm and three replicates of L*er nrpd1*-/- female x Col-0 male (referred to as mat *nrpd1*+/-) endosperm. Because the endosperm is triploid, in these comparisons there are 3 (wild-type), 2 (pat *nrpd1*+/-), 1 (mat *nrpd1*+/-) and 0 (*nrpd1*-/-) functional *NRPD1* alleles in the endosperm. However, *NRPD1* is a paternally expressed imprinted gene in wild-type L*er* x Col endosperm and the single paternal allele contributes 62% of the *NRPD1* transcript whereas 38% comes from the two maternal alleles (Pignatta et al., 2014). Consistent with paternal allele bias in *NPRD1* expression, mRNA-Seq data shows that *NRPD1* is expressed at 42% of wild-type levels in pat *nrpd1*+/- and at 91% of wild-type levels in mat *nrpd1*+/- (Supplementary Table 6).

We found that the presence of functional *NRPD1* inherited from either parent is sufficient for the biogenesis of nearly wild-type levels of 21-24 nt sRNAs in endosperm (Fig. 1A, Fig. S3). However, although the overall sRNA population in the heterozygotes was similar to the wild-type (Fig. 1A), loss of maternal and paternal *NRPD1* had distinct impacts on sRNA at individual loci (Fig. 1B-F, Fig. S3, Table S2-3). We identified genes and transposable element (TE) insertions that displayed at least a two-fold change in the abundance of sRNAs in mat or pat *nrpd1*+/- compared to the wild-type (Fig. 1B-F, Fig. S3, Tables S2-3). Loss of paternal *NRPD1* caused relatively small fold-change reductions in 21-24 nt Pol IV sRNAs at a handful of loci, while loss of maternal *NRPD1* had slightly greater yet limited impact (Fig. 1B-F, Fig. S3, Table S2-3). For genic loci with *NRPD1*-dependent 24 nt sRNAs, 2% (327 genes) had significantly lower abundance in mat *nrpd1*+/- compared to wild-type; in contrast 0.3% (60 genes) were significantly lower in pat *nrpd1*+- (Fig. 1B). For TE loci with *NRPD1*-dependent 24 nt sRNAs, 2.8% (545 TE insertions) and 1.35% (261 TE insertions) exhibited significantly lower abundance in mat and pat nrpd1+/-, respectively (Fig. 1B). Few of the loci with reduced sRNAs were shared between the reciprocal heterozygotes – of 327 24nt sRNA-expressing genic loci that were reduced by more than two-fold in mat *nrpd1*+/-, only 22 were also reduced by two-fold in pat *nrpd1*+/- (Fig. 1F). Moreover, there was no quantitative or correlative relationship between loci affected in mat *nrpd1*+/- and pat *nprd1*+/- (Fig. 1F). Thus, the vast majority of sRNA-producing loci in endosperm only require at least one functional copy of *NRPD1* after fertilization.

**Fig. 1:**
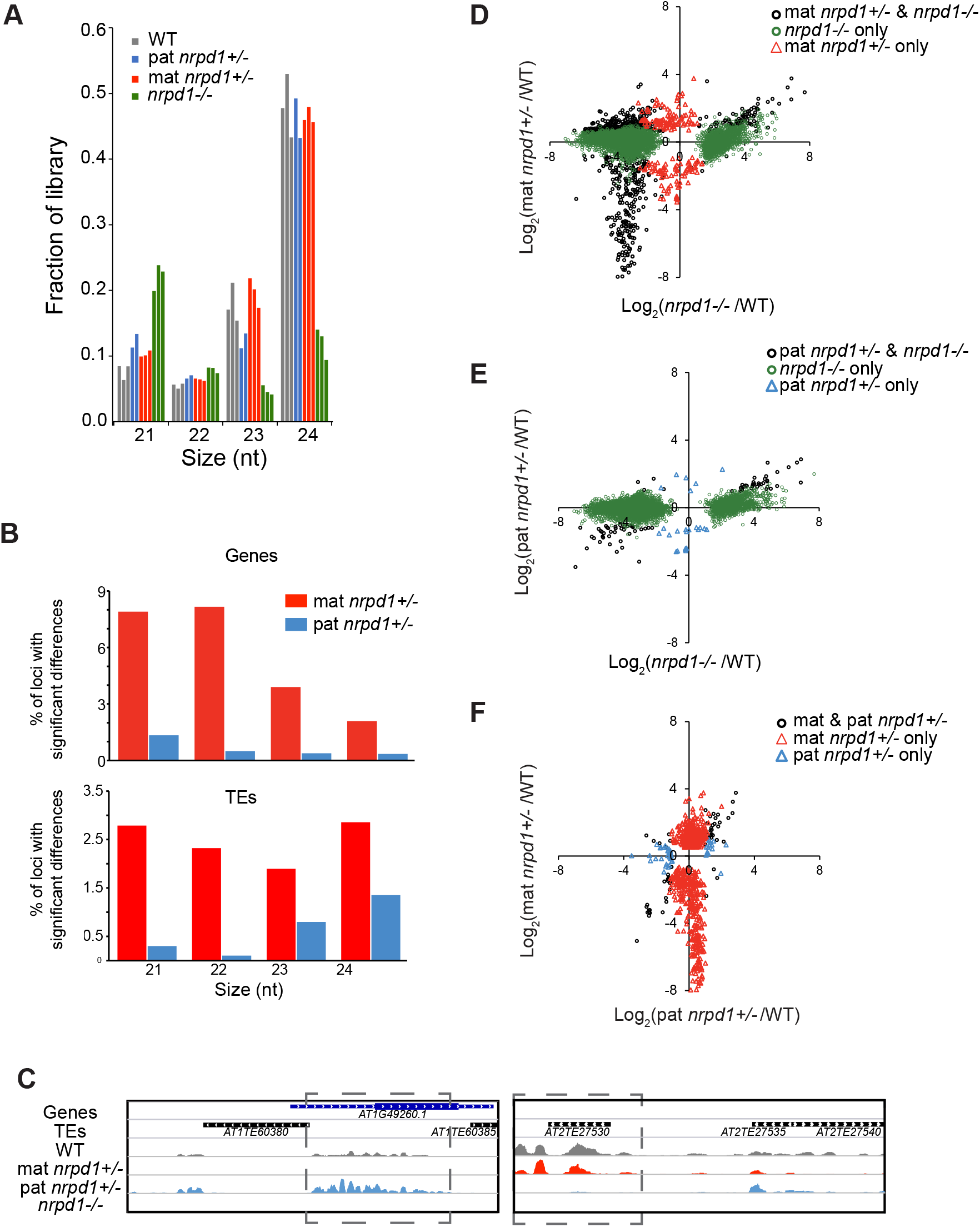
Impact of loss of maternal, paternal, or both copies of *NRPD1* on endosperm small RNAs. (**A**) Loss of maternal or paternal *NRPD1* does not substantially alter the endosperm small RNA pool. Fraction of aligned small RNA reads in each size class in the indicated genotypes. (**B**) Examination of 21-24 nt sRNAs over genes or TEs shows that inheriting a mutant maternal *nrpd1* allele has a larger impact than inheriting a mutant paternal *nrpd1* allele. Percent of loci showing at least a two-fold reduction in sRNA abudance and padj <0.05 according to DESeq2 are indicated in red (mat *nrpd1*+/-) or blue (pat *nrpd1*+/-). Genes and TEs included in this tally have a normalized wild-type read count of five or higher. (**C**) Snapshots of loci with Pol IV-dependent 24 nt sRNAs that show a specific loss of small RNAs in mat (left) or pat (right) *nrpd1*+/- endosperm. (**D-F**) Comparisons of genic 24 nt sRNAs upon loss of maternal, paternal, or both copies of *NRPD1*. Fold change as calculated by DESeq2. Only significant changes (q<0.05) are plotted.

### Evaluating memory of parental Pol IV activity and endosperm sRNA production

The absence of dramatic differences in sRNAs in heterozygotes could indicate that the alleles inherited from both the wild-type and the *nrpd1*-/- parent produce a wild-type level of sRNAs after fertilization. This result would be expected for a recessive mutation without parental effects. However, it is known that Pol IV activity at some loci requires prior Pol IV activity (Jingwen Li et al., 2020). Under such a scenario, Pol IV activity in the parents before fertilization might be necessary for sRNA production from that parent’s allele in the endosperm after fertilization. Thus, the observed lack of differences in sRNA production at most loci in heterozygous *nrpd1* endosperm (Fig. 1) could be explained by an upregulation of sRNA production from the alleles inherited from the wild-type parent (i.e. sRNAs are upregulated from paternal alleles in mat *nrpd1*+/- and maternal allele sRNAs are upregulated in pat *nrpd1*+/- endosperm). To distinguish between these possibilities, we used the SNPs between Col-0 and L*er* to identify the allelic origins of small RNAs in WT and heterozygous endosperm. We first confirmed prior observations that Pol IV sRNAs are biallelically expressed at most loci in endosperm and predominantly expressed from one parental allele, or imprinted, at several hundred others (Erdmann et al., 2017). Examining sRNAs at genes and TEs, we found that both bi-allelically expressed 21 and 24 nt sRNA loci (defined as between 20% and 80% of sRNAs from maternally-inherited alleles) and those predominantly expressed from one parental allele (>80% or <20% maternal) were Pol IV-dependent (i.e. their accumulation was significantly reduced in *nrpd1*-/- endosperm) (Fig. S4).

To test if sRNA production from alleles inherited from wild-type parents compensated for alleles inherited from an *nrpd1*-/- parent, we first assessed several thousand loci that were not significantly mis-regulated in *nrpd1*+/- endosperm. Overall, there were similar contributions from maternal and paternal alleles in mat and pat *nrpd1* heterozygotes compared to wild-type endosperm (Fig. 2A). This suggests that by 7 DAP (days after pollination), at most loci in the endosperm, sRNAs are produced from both maternal and paternal alleles regardless of whether the alleles were inherited from a wild-type parent or an *nrpd1*-/- parent. However, we found that imprinted sRNA regions (ISRs) (113.1KB maternally imprinted and 1215.6KB paternally imprinted regions overlapping both genic and TE loci.) (Erdmann et al., 2017) were impacted by loss of parental Pol IV (Fig 2C-F). 179 of 206 ISRs where expression is maternally biased in WT showed reduced 24 nt sRNAs in mat *nrpd1*+/- (Fig. 2C). ISR loci have been filtered to remove regions that are also enriched for seed coat sRNAs (Erdmann et al., 2017) and thus preclude analytical artifacts that may arise due to maternal tissue contamination or due to any potential sRNA movement. On the other hand, only a small subset (74 of 2405 ISRs) of paternally biased ISRs produced fewer sRNAs in pat *nrpd1*+/- and slightly more sRNAs in mat nrpd1+/-. We also note that maternally biased regions in wild-type showed slightly elevated production of sRNAs in pat *nrpd1*+/- endosperm and paternally biased regions in wild-type show slightly elevated levels of sRNAs in mat *nrpd1*+/- endosperm (Fig. 2D). Examination of the allelic origins of sRNAs at genes and TEs are also consistent with the ISR analysis. In a parallel analysis, we found that small RNA loci showing dramatic reductions in abundance in mat *nrpd1*+/- tended to be maternally biased in wild-type endosperm (>80% of sRNAs from the maternally-inherited alleles) (Fig. 2B, leftmost column). Similarly, in pat *nrpd1*+/-, paternally biased small RNA (<20% sRNAs from the maternally-inherited alleles) loci were more impacted (Fig. 2B, rightmost column).

**Figure 2:**
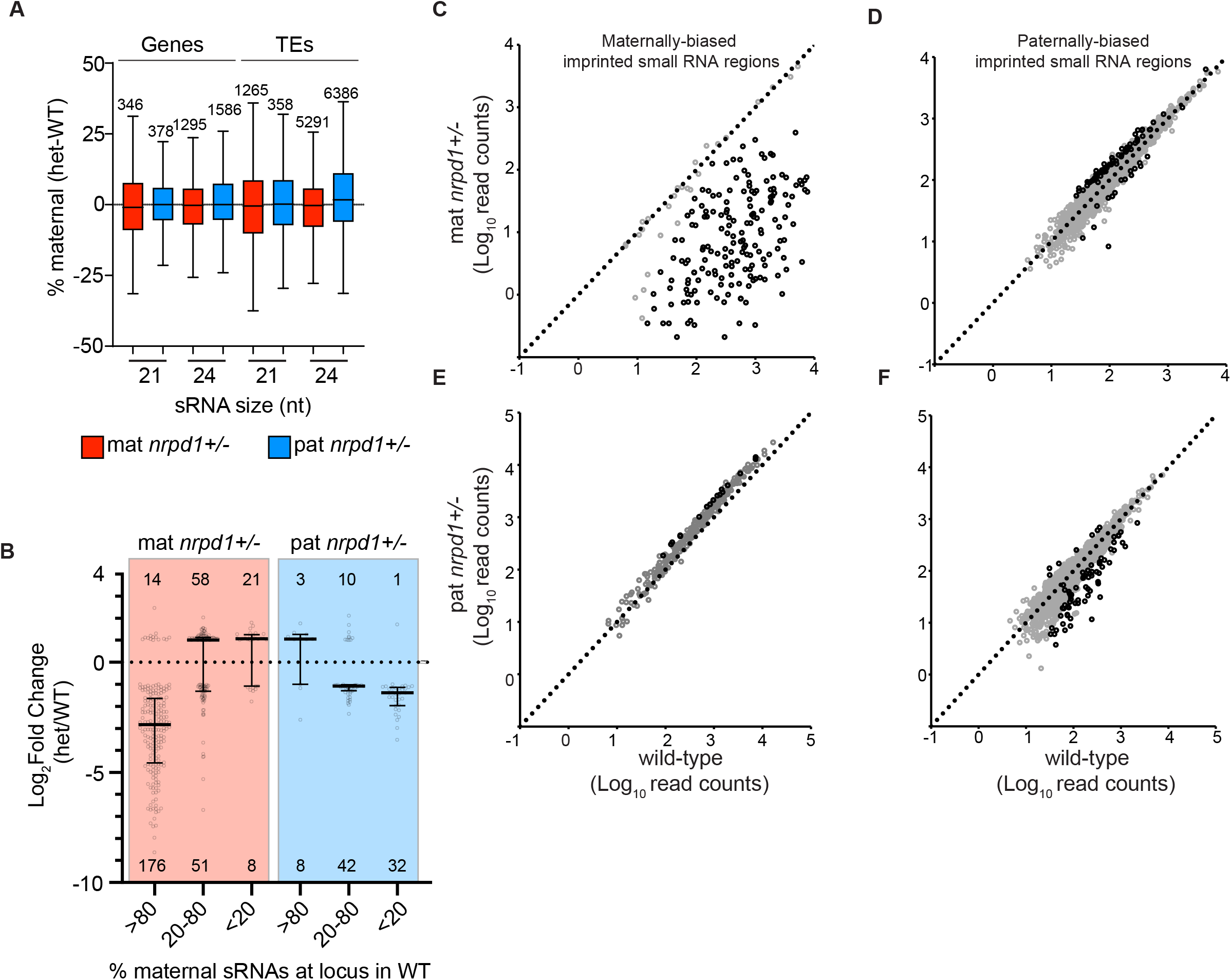
Effects of loss of maternal or paternal Pol IV on the allelic origin of small RNAs. **(A)** Tukey plot shows no difference in allelic origin of genic and TE small RNAs between heterozygotes and wild-type. Loci plotted here show similar abundances in wild-type and heterozygotes and have a sum of at least ten allele-specific reads in three wild-type replicates and in heterozygotes. **(B)** Loci with reduced sRNAs in mat *nrpd1*+/- or pat *nrpd1*+/- exhibit maternally- or paternally-biased sRNAs in WT. Genes and TE showing differential abundance of 24 nt sRNAs in *nrpd1* heterozygotes were grouped into bins by the % of sRNAs produced from the maternal alleles of that locus in WT. Fold-change was calculated by DESeq2.Tukey plot represents fold-change in each group. Circles show fold-change at individual loci. Numbers below and above plot are total number of loci having significantly lower and higher abundance of 24nt sRNAs in *nrpd1*+/- relative to the wild-type. (**C-F**) Loss of maternal *NRPD1* leads to a reduction in the abundance of sRNAs from maternally-biased ISRs (imprinted small RNA region) (C) and gain of sRNAs from a subset of paternally-biased ISRs (D). Loss of paternal *NRPD1* has a negligible impact on maternally-biased ISRs (E) and a relatively minor impact on paternally-biased ISRs (F). Col-L*er* imprinted sRNA regions used here were defined in Erdmann *et al* (2017). These regions have less expression in seed coat relative to endosperm. To identify regions with changes in small RNA abundance, read counts were calculated over sliding windows of 300bp with 200bp overlap. Windows with differential abundance were identified using DESeq2. Windows overlapping an ISR were identified using bedtools intersect. Overlapping windows were merged using bedtools merge and the median read-count for each set of merged windows was plotted. Windows with and without significant differences in abundance are represented by black and grey circles.

In summary, these results indicate that imprinted sRNA loci in endosperm are dependent on Pol IV activity in the parents and are not established *de novo* post-fertilization. Notably, these sites of Pol IV action are by definition distinct between maternal and paternal parents.

### Maternal and paternal RNA Pol IV have antagonistic impacts on gene expression

We previously identified several hundred genes mis-expressed in *nrpd1*-/- endosperm. To test for maternal or paternal effects on endosperm gene expression, we performed mRNA-seq in three replicates each of mat *nrpd1*+/- and pat *nrpd1*+/-, along with appropriate wild-type controls and homozygous mutant *nrpd1* endosperm (Table S1). Examination of these datasets using a tissue-specific gene expression tool showed no indication of contamination with seed coat tissue (Figure S5). Differential expression analyses identified 1791 genes whose transcripts were more abundant and 1455 that were less abundant in *nrpd1*-/- compared to wildtype endosperm (Fig. 3; Table S6). Almost 50% of these genes (1599) were similarly mis-regulated in mat *nrpd1*+/- (Fig. 3A,B), along with 2998 additional genes. In contrast, very few genes (90) changed in expression in pat *nrpd1*+/- compared to the wild type (Fig. 3A, B). In addition to the difference in the size of the effect, loss of maternal or paternal Pol IV altered the expression of different classes of genes. Panther over-representation tests (Mi & Thomas, 2009) indicated that in mat *nrpd1*+/-, down-regulated genes were enriched for functions in the cell-cycle, whereas up-regulated genes were enriched for functions in photosynthesis, stress response, and abscisic acid signaling. In pat *nrpd1*+/-, up-regulated genes were enriched for functions in heat stress response, while down-regulated genes were enriched for functions in responses to fungi. The expression of imprinted genes is known to be regulated epigenetically in endosperm. 15 paternally expressed and 45 maternally expressed imprinted genes were mis-regulated in mat *nrpd1*+/- while two maternally expressed imprinted genes but no paternally expressed imprinted genes were mis-regulated in pat *nrpd1*+/- (Table S6).

**Figure 3.**
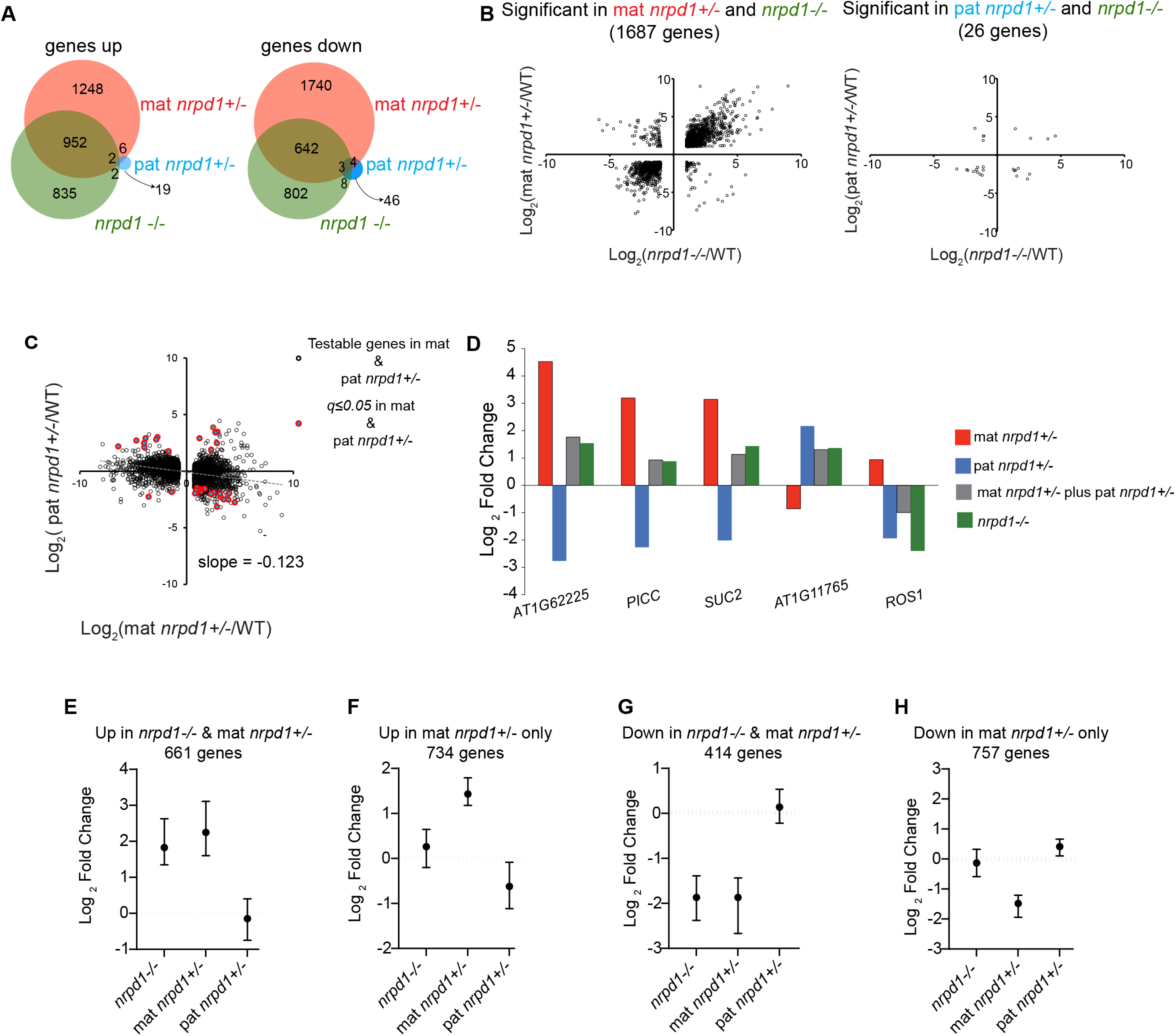
Maternally and paternally acting Pol IV have antagonistic effects on endosperm gene expression. **(A)** Venn diagrams showing overlap of genes with increased and decreased expression in comparison to wild-type endosperm for the indicated genotypes. **(B)** Scatter plots of genes that are all significantly different (*q*≤*0.05*, Log_2_(Fold Change) ≥1 or ≤ −1) between wild-type and indicated mutants. Fold-change calculated using Cuffdiff. **(C)** Inverse correlation between changes in gene expression in mat and pat *nrpd1*+/- relative to WT. Slope was calculated for all testable genes in comparisons of mat and pat *nrpd1*+/- relative to WT. Genes significantly mis-regulated in both mat and pat *nrpd1*+/- are colored circles. **(D)** Examples of genes that are antagonistically regulated by Pol IV. Grey bars represent mathematical sum of effects observed in mat and pat *nrpd1*+/-. **(E)** Genes up-regulated at least two-fold in both *nrpd1*-/- and mat *nrpd1*+/- do not exhibit mis-regulation in pat *nrpd1*+/-. **(F)** Genes up-regulated only in mat *nrpd1*+/- but not *nrpd1*-/- have decreased expression in pat *nrpd1*+/-. **(G)** Genes down-regulated in both *nrpd1*-/- and mat *nrpd1*+/- are not mis-regulated in pat *nrpd1*+/-. **(H)** Genes down-regulated only in mat *nrpd1*+/- but not in *nrpd1*-/- are overall slightly increased in expression in pat *nrpd1*+/-. Plots E-H show median and inter-quartile range for log_2_ fold change in mutant/WT. Fold-change were calculated by Cuffdiff.

Differential expression of a gene between wild-type and *nrpd1*-/- could represent: 1) maternal and paternal effects arising from the loss of *NRPD1* in parents, 2) zygotic effects arising from epistatic interactions between mat *nrpd1*- and pat *nrpd1*-, 3) effects from the loss of all *NRPD1* in the endosperm, or 4) the sum of all three effects. As this study does not examine the effect of knocking-down *NRPD1* specifically in the endosperm, we can only detect parental effects. Curiously, 2988 genes mis-regulated in mat *nrpd1*+/- were not mis-regulated in *nrpd1*-/- endosperm (Fig. 3A). We hypothesized that genic mis-regulation found exclusively in mat *nrpd1*+/- (but not *nrpd1*-/-) was caused by separate transcriptional effects of maternal and paternal *nrpd1* that were obscured in null mutants. To test this hypothesis, we compared gene expression between mat and pat *nrpd1*+/- (Fig. 3C). We found that 51/90 genes mis-regulated in pat *nrpd1*+/- endosperm were also mis-regulated in mat *nrpd1*+/- endosperm (red circles in Fig. 3C). However, 36 of these 51 genes changed expression in the opposite direction. For example, expression of the gene *SUC2* decreased about four-fold in pat *nrpd1* +/- endosperm and increased about eight-fold in mat *nrpd1*+/- endosperm (Fig. 3D). If *NRPD1* loss has no endospermic (zygotic) effect on the expression of these genes, then the mis-regulation observed in *nrpd1*-/- endosperm would be the sum of the parental effects. Indeed, the change in abundance of these genes in *nrpd1*-/- endosperm is close to that predicted by an additive, antagonistic parental effect (compare gray and green bars in Fig. 3D). *SUC2* transcript abundance in *nrpd1*-/- changes by 2.7-fold compared to the predicted 2.18-fold change, and other genes show similar effects (Fig. 3D). While the expression of these particular genes showed large effects in both heterozygotes, most genes mis-regulated in mat *nrpd1*+/- did not show a significant change (>2-fold difference in transcript abundance) in pat *nrpd1*+/- endosperm. We therefore hypothesized that mis-regulation of genes in mat *nrpd1*+/- but not *nrpd1*-/- endosperm was due to a small antagonistic effect arising from the loss of pat *NRPD1* in *nrpd1*-/-. Indeed, genes mis-regulated in either mat *nrpd1*+/- or pat *nrpd1*+/- (Fig. 3C) are overall negatively correlated (slope = −0.123). To test this hypothesis further, we evaluated the expression of genes in pat *nrpd1*+/- endosperm that were either mis-regulated in both mat *nrpd1*+/- and *nrpd1*-/- or only in mat *nrpd1* +/-. Transcripts that significantly increased exclusively in mat *nrpd1*+/- had slightly decreased expression in pat *nrpd1*+/- endosperm (Fig. 3F). In contrast, genes that were significantly upregulated in both mat *nrpd1*+/- and *nrpd1*-/- were not affected in pat *nrpd1*+/- (Fig. 3E). Similarly, genes that showed a significant reduction in abundance only in mat *nrpd1*+/- were slightly higher expressed in pat *nrpd1*+/- endosperm (Fig. 3H) while genes with reduced abundance in both mat *nrpd1*+/- and *nrpd1*-/- were not affected in pat *nrpd1*+/- (Fig. 3G). These results are consistent with an antagonistic parent-of-origin effect model for the impact of Pol IV on endosperm transcriptomes. Although the antagonistic effect at most genes is less than the commonly used two-fold threshold difference for a significant change in gene expression, it is similar in magnitude to dosage compensation effects in other systems such as that observed for the fourth chromosome in *D. melanogaster* and for genes mediating genetic compensation in zebrafish (El-Brolosy et al., 2019; Johansson et al., 2012).

### Evaluating possible mechanisms of Pol IV’s impact on gene expression

How does Pol IV have parent-of-origin effects on gene expression in the endosperm after fertilization? Pol IV effects could be direct or indirect at the affected loci. One possibility is that Pol IV modulates gene expression via the proposed post-transcriptional gene silencing (mRNA cleavage) or translational inhibition by 21-22nt Pol IV-dependent small RNAs (Jullien et al., 2020; Panda et al., 2020). Or, Pol IV-dependent RNA-directed DNA methylation over genic sequences or linked gene regulatory element might repress transcription in the wild type. Alternatively, Pol IV could impact many genes in *trans* by regulating the expression of chromatin proteins, like the known target *ROS1* (a DNA demethylase) (Williams et al., 2015), transcription factors (Kirkbride et al., 2019; Xu et al., 2015), or by broadly influencing genome organization, which in turn affects gene expression (Rowley et al., 2017; Zhong et al., 2021). To estimate the contribution of *cis* and *trans* effects and identify potential cis-regulatory targets of Pol IV that could drive wide-spread *trans*-effects, we analyzed the congruence of small RNAs, DNA methylation, mRNA cleavage patterns, and allele-specific changes driving gene expression changes in wild-type and mutant endosperm.

#### Assessing potential mRNA cleavage by Pol IV-dependent sRNAs in endosperm

Pol IV dependent genic sRNAs are proposed to regulate gene expression by cleaving mRNA (Panda et al., 2020) and we previously demonstrated that endosperm has greater accumulation of genic sRNAs than other tissues (Erdmann *et al*, 2017). To test if such cleavage events contribute to endosperm Pol IV-dependent transcript abundance, we first identified candidate genes that exhibited significantly increased (≥ 2-fold) mRNA abundance and significantly decreased 21, 22, or 24 nt sRNA abundance (≥2-fold) in *nrpd1*-/- endosperm (Fig. S6A-C). This analysis suggested that at least 305 genes or 16% of the genes that increase in expression in *nrpd1*-/- endosperm were associated with Pol IV dependent sRNAs. To directly assay if these genic sRNAs drive mRNA cleavage at levels sufficient to alter transcript abundance at specific loci, we mapped the 5’ ends of mRNA from wild-type and *nrpd1*-/- endosperm mRNA using NanoPARE sequencing (Schon et al., 2018). NanoPARE maps both the 5’ ends of primary transcripts and those that result from mRNA cleavage. We confirmed that NanoPARE sequencing was working for us by identifying transcriptional start sites as well as internal cleavage sites for known miRNA targets (Fig. S6E). We found that almost all genes exhibiting increased mRNA abundance also exhibited increased 5’ ends at transcriptional start sites, but did not have reduced 5’ ends internal to gene (Fig. S6D). This was confirmed by visual observation of individual loci (Fig. S6E). This suggests that the increase in the transcript abundance of these genes in *nrpd1*-/- is not caused by reduced mRNA cleavage. Only five genes – *PERK8*, *GLP2A*, *ETTIN*, *AAD3* and *AT2G45245* – exhibited reduced cleavage at a few sites in *nrpd1*-/- endosperm. This minimal effect is contrary to the expectation that candidate genes described above should have reduced cleavage and suggests that small RNA mediated post-transcriptional gene silencing is not a key mechanism for Pol IV to control endosperm gene expression. We therefore did not test if mRNA cleavage is impaired in *nrpd1*+/- heterozygotes.

#### Assessing correspondence between sRNA and mRNA changes

Only a minority of the genes that have altered expression in *nrpd1* endosperm have associated changes in sRNAs within those same genes (Fig. S6A-C). However, Pol IV sRNAs may also act at sites proximal to a gene to regulate it. We assessed the distance between mis-expressed genes and altered sRNAs in homozygous and heterozygous *nrpd1* mutant endosperm. We found that 9.2%, 11.7% and 3.3% of mis-regulated genes are within 1 kb of a site that loses sRNAs in *nrpd1*-/-, mat *nrpd1*+/-, and pat *nrpd1*+/- endosperm, respectively (Table S7). To obtain a genome-wide perspective not focused on arbitrary distance cutoffs, we used the relative distance metric to test if genomic regions losing 24nt sRNAs were associated with mis-regulated genes. The relative distance metric describes the spatial correlation between sRNA intervals and mis-regulated genes, compared to mis-regulated genes and random intervals (Favorov et al., 2012). This analysis found no enrichment in the association between Pol IV dependent sRNAs and mis-regulated genes in any of the genotypes (Fig. S6F).

#### Assessing correspondence between DNA methylation and mRNA changes

The relevant molecular function of RNA Pol IV with regard to gene expression is typically assumed to be its role in RdDM. To identify potential examples of DNA methylation mediating Pol IV’s impact on genes, we performed bisulfite sequencing of WT, mat *nrpd1*+/-, and pat *nrpd1*+/- endosperm DNA. We evaluated wild-type DNA methylation at individual cytosines within Pol IV sRNA-producing genes that were mis-regulated in *nrpd1*-/-, mis-regulated genes that showed increased sRNAs in *nrpd1*-/- (Pol IV independent), and five control sets of genes that showed no change in sRNA abundance upon loss of Pol IV. We found that most cytosines were not methylated (median is near zero) in the genes we examined (Fig. S7A). This suggests that Pol IV small RNAs do not generally target DNA methylation at genes. However, some mis-regulated genes with Pol IV dependent sRNAs had higher CG methylation in wild-type endosperm (Fig. S7A). These mis-regulated genes with Pol IV dependent sRNAs were also likely to be longer (Fig. S7B). Longer genes have higher small RNA read counts (Fig. S7C) and thus differences in these genes are more likely be called as statistically significant by DESeq2 (Oshlack & Wakefield, 2009). Longer genes also tend to have higher CG methylation (Takuno & Gaut, 2012; Zilberman et al., 2007). We therefore argue that this increased CG methylation is an analytical artifact. Our results suggest that Pol IV dependent genic sRNAs do not regulate endosperm gene expression by directing genic DNA methylation, consistent with our previous findings (Erdmann et al., 2017).

We also tested if changes in DNA methylation brought about by loss of parental Pol IV could explain changes in gene expression. Overall, loss of parental Pol IV had only minor effects on DNA methylation (Table S5). Loss of Pol IV activity primarily reduces asymmetric CHH methylation (Stroud et al., 2013). Comparison of mat *nrpd1*+/- and pat *nrpd1*+/- CHH methylation with wild-type endosperm identified 2234 and 2056 DMRs (covering 812.7 Kb and 759.9Kb, respectively) with 50% hypomethylated in mat *nrpd1*+/- and 54.8% hypomethylated in pat *nrpd 1*+/- (Table S5). Consistent with the parent-of-origin effects described for mRNAs and sRNAs, we made three observations that suggest that mat and pat Pol IV activity have distinct impacts on the endosperm methylome. First, only 50% of CHH DMRs are shared between the two heterozygous genotypes. Second, regions where sRNA accumulation is dependent on paternal inheritance of a wild-type *NRPD1* allele have higher CHH methylation in wild-type endosperm than regions where sRNAs are dependent on maternal *NRPD1* (Fig. S7D). This pattern is consistent with our previous finding that maternally-biased small RNAs are often not associated with methylated DNA in wild-type endosperm (Erdmann et al., 2017). Third, an examination of regions with at least 10% CHH methylation in wild-type endosperm shows that loss of paternal *NRPD1* had a more substantial impact on endosperm CHH methylation than loss of maternal *NRPD1* (Fig. S7E).

Symmetric CG and CHG methylation are typically less affected by loss of *NRPD1* because other mechanisms exist to maintain this type of methylation. Comparison of CHG methylation between wild-type and either heterozygote identified fewer than 100 DMRs and CHG methylation was not investigated further. Both mat and pat *nrpd1*+/- endosperm exhibited changes in CG methylation compared to the wild type (Table S5). In mat *nrpd1*+/- endosperm, 48.5% of DMRs (of a total 600 DMRs spanning 207 kb) were hypomethylated relative to wild-type while in pat *nrpd1*+/- 60% of DMRs (of a total 707 DMRs spanning 258 KB) were hypomethylated relative to wild-type. Further, we found that few of the sites hypo- or hyper-methylated in the CG context in mat *nrpd1*+/- were shared with those changing methylation state in pat *nrpd1*+/- (Table S5).

We used the DMRs in the analysis described above to assess their impact on gene expression. In mat *nrpd1*+/- endosperm, 2.6% and 3.4% of total mis-regulated genes are within one kb of assayable regions with less or more CHH methylation in mat *nrpd1*+/- endosperm (Table S7). In addition, two genes and one gene are within one kb of a region that has higher and lower CHH methylation in pat *nrpd1*+/-. One gene is associated with increased CG methylation in pat *nrpd1*+/-. We also used relative distance analysis to see if mat *nrpd1*+/- mis-regulated genes are more likely to be associated with DNA methylation changes (there are too few genes associated with DNA methylation changes in pat *nrpd1*+/- to perform this analysis). Consistent with previous analyses, we find no clear relationship between DNA methylation changes and gene expression changes in the mat *nrpd1*+/- endosperm (Fig. S7F).

#### *Allelic analysis of mis-regulated genes to identify* cis *or* trans *effects of Pol IV*

One method to assess whether Pol IV’s impacts on gene expression are predominantly *cis* or *trans* acting is to compare the allelic origins of mRNA in wild-type and *nrpd1*+/- endosperm. If a gene’s mRNA abundance in the endosperm is determined by the activity of Pol IV in *cis* either in the gametophyte or sporophyte, then the gene would be primarily mis-regulated from the allele inherited from a parent lacking Pol IV. Thus, in mat *nrpd1*+/- endosperm, mis-regulation of such genes would be driven predominantly by changes in expression from maternal alleles whereas genes expression differences in pat *nrpd1*+/- endosperm would be driven by changes in expression from paternal alleles. In contrast, the predominance of *trans* effects would be indicated by both parental alleles contributing to the changes in the abundance of transcript levels at most genes. We utilized SNPs between Col-0 (paternal) and L*er* (maternal) genomes to identify allele-specific mRNA-seq reads. We evaluated the contributions of each parent’s alleles in the endosperm for 2372 mis-regulated genes that had at least ten allele-specific reads in wild-type and *nrpd1*+/-. For the majority of genes, mis-regulation in mat *nrpd1*+/- was driven by effects on expression of both maternal and paternal alleles, with some notable exceptions (Fig. S8A). For example, increased expression of *DOG1* in mat *nrpd1*+/- was primarily driven by increased expression from maternal alleles (Fig. S8B). *AT4G12870* was repressed in mat *nrpd1*+/- primarily due to a loss of maternal allele expression (Fig. S8B). In contrast, expression of *SAC2* was primarily repressed in mat *nrpd1*+/- because of decreased expression from the paternal allele (Fig. S8B). Overall, both maternal and paternal alleles made equal contributions to genic mis-regulation in the mat *nrpd1*+/- endosperm. 4.7% of down-regulated genes and 5.3% of up-regulated genes showed at least a 20% increase or decrease in maternal allele contribution. This was roughly similar to the contribution of paternal alleles to mis-regulation in mat *nrpd1*+/-. 4.4% of down-regulated genes and 5.3% of up-regulated showed at least a 20% change in paternal allele contribution (Fig. S8A). In pat *nrpd1*+/-, only 8% of down-regulated genes had lower contribution of paternal alleles while both alleles contributed to up-regulation (Fig.S8A). Overall, these results suggest that parental Pol IV’s impact on gene expression is largely due to *trans* effects.

In summary, our analyses test and dismiss several *cis*-regulatory mechanisms for how Pol IV may mediate parent-of-origin gene expression effects on the endosperm. We also individually examined DNA methylation and sRNAs at genes showing antagonistic regulation by maternally and paternally acting Pol IV and found no evidence for a role for sRNAs and DNA methylation in their regulation. These results lead us to conclude that parent-of-origin effects and the antagonistic effects that we observe are likely the result of *trans*-acting effects of parental Pol IV activity.

## Discussion

We demonstrate that Pol IV activity in the father promotes seed abortion in response to extra paternal genomes, whereas Pol IV activity in the mother promotes seed viability in these conditions. Previous observations of sRNA or mRNA at individual genes in diploid endosperm showed that Pol IV function in the mother and the father have different effects on the endosperm (Kirkbride et al., 2019; Vu et al., 2013). These findings suggested that Pol IV has differing, and perhaps even opposing, roles in maternal and paternal parents. In this study, we characterized the effect of maternal and paternal Pol IV activity on the endosperm through genome-wide analyses of transcription, mRNA cleavage, small RNAs, and DNA methylation in balanced endosperm. Our molecular data demonstrate that Pol IV activity in the mother and father have parent-of-origin effects on the endosperm, a subset of which are antagonistic. We found that one parent’s copy of *NRPD1* is sufficient for the production of Pol IV-dependent sRNAs at most loci, with a small number of largely non-overlapping loci losing sRNAs upon loss of maternal or paternal *NRPD1*. Pol IV activity in the mother and father also have distinct impacts on the DNA methylation landscape in the endosperm. Endosperm with a paternally inherited *nrpd1* mutation had lower DNA methylation compared with endosperm where the *nrpd1* mutation was maternally inherited. Finally, an interrogation of gene expression shows that loss of maternal Pol IV leads to significant mis-regulation of several hundred genes while loss of paternal Pol IV leads to mis-regulation of only several dozen. A key finding of our study is that genes that are mis-regulated upon loss of maternal *NRPD1* are affected in an opposite manner upon loss of paternal *NRPD1*. Together, our results suggest that maternal and paternal Pol IV are genetically antagonistic and that the major effect on transcription observed in heterozygotes is established before fertilization. These observations are important for understanding both Pol IV’s role in reproduction and the genetic architecture underlying parental control of offspring development.

### Pol IV, conflict, and the genetic architecture of parental control

Parental conflict theory predicts that in viviparous species, mother and father have antagonistic effects on regulating resource allocation and associated gene expression in offspring. In practice, such effects are difficult to detect and have been infrequently described. Antagonistic parental effects are likely to be balanced in the individuals within an inbred population (like Arabidopsis) and are thus unobservable except in mutants or in hybrids where maternal and paternal effects are out of balance. When homozygous mutants are examined, these effects may be missed because they do not cause dramatic developmental phenotypes or because simultaneous loss of antagonistic maternal and paternal effects effectively cancels one another out. Thus, reciprocal heterozygotes need to be examined to detect antagonistic parent-of-origin effects. A close examination of our data provides insights into the genetic architecture mediating parental control of offspring development.

A key feature of the regulatory infrastructure that mediates parent-of-origin specific effects on zygotic gene expression is that maternal and paternal alleles need to be distinguished from each other in the zygote (in this case, endosperm is the relevant zygote). In *A. thaliana* endosperm, at many loci maternally inherited alleles are DNA demethylated and marked with H3K27 methylation by Polycomb Repressive Complex2 (PRC2), while paternally inherited alleles remain DNA methylated and have reduced H3K27me3 (Borg et al., 2020; Moreno-Romero et al., 2016; Pignatta et al., 2014). Maternal inheritance of mutations in the PRC2 sub-units *MEA*, *FIE*, *FIS2* and *MSI1* leads to endosperm defects and seed abortion (Chaudhury et al., 1997; Grossniklaus, 1998; Kohler, 2003; Ohad et al., 1996). Similarly, inheritance of maternal mutations in the DNA demethylase *DME* increases DNA methylation on endosperm maternal alleles and causes seed abortion (Choi et al., 2002). Paternal inheritance of mutations in these genes have no reported effect on endosperm development or gene expression. These results thus argued that the solution to the problem of distinguishing parental alleles from one another after fertilization was to mark maternal and paternal chromosomes with distinct epigenetic modifications. However, this model may not explain all parent-of-origin effects on gene expression, particularly outside of imprinted genes. Our study provides evidence for a distinct model in which the same epigenetic regulator – Pol IV – can mediate both maternal and paternal effects. The only other example of a gene with seemingly antagonistic effects on seeds is the maintenance methyltransferase *MET1*, whose mutation has opposing effects on seed size when inherited maternally or paternally, although the molecular basis of this phenotype is unknown (Xiao et al., 2006).

How does Pol IV in the mother and the father have distinct impacts after fertilization? Pol IV targets can be tissue or developmental stage-specific (Grover et al., 2020) and thus Pol IV may target different genomic regions during male and female gametogenesis. Pol IV could act pre-or post-meiotically in the parental sporophyte (diploid phase of the life cycle), in the gametophyte (haploid phase of life cycle), or post-fertilization in the maternal sporophyte. RT-PCR based examination of dissected synergids and central cells did not detect *NRPD1* transcripts (Vu et al., 2013). This suggests that on the maternal side Pol IV influences endosperm gene expression by acting in the maternal sporophyte or in the female gametophyte prior to central cell formation. Alternatively, Pol IV could act in the maternal sporophytic integuments/seed coat after fertilization, when the endosperm is developing. One potential mechanism for this would be through Pol IV-dependent sRNAs moving from the seed coat to the endosperm (Grover et al., 2020; Kirkbride et al., 2019). However, examination of the levels of total Pol IV-dependent sRNAs, allele-specific data, and imprinted sRNA regions suggests that the potential influence of seed coat Pol IV function on endosperm expression would likely be independent of sRNA transfer. This conclusion is consistent with previous observations that endosperm and seed coat have distinct sRNA profiles (Erdmann et al., 2017).

We have shown that parental Pol IV activity is dispensable for guiding endosperm sRNA production at most loci, with the exception of imprinted sRNA regions, but that parental Pol IV activity plays an important role in guiding endosperm gene expression. The molecular nature of this memory is unknown, and at present we can only speculate. Data from paternal excess interploidy crosses suggests that the molecular identity of Pol IV memory may differ between the maternal and paternal parents. In the father, the genes required for sRNA production (*NRPD1*, *RDR2* and *DCL3*) and the genes required for downstream DNA methylation (*NRPE1/Pol V* and *DRM2*) are both essential to promote paternal excess seed abortion (Satyaki & Gehring, 2019). In contrast, in the mother, genes required for sRNA production but not for DNA methylation promote paternal excess seed viability (Satyaki & Gehring, 2019). This suggests that DNA methylation or another downstream chromatin mark directed by Pol IV-dependent sRNAs could be the identity of paternally-inherited memory, but is unlikely to be the molecular identity of maternally-inherited memory. What would be the nature of maternal DNA methylation-independent memory? Pol IV, like other RNA polymerases (Studitsky et al., 2004), could act as a chromatin remodeler. Or, Pol IV could direct a chromatin modification, produce sRNAs that post-transcriptionally control genes, or control the expression of genes whose products are deposited in the gametes, which in turn sets up a memory to direct gene expression programs in the endosperm after fertilization.

How might we interpret Pol IV’s parent-of-origin effects in terms of conflicts between parents? Studies on how resource allocation conflicts between parents impact gene expression have thus far been focused on imprinted genes. However, a handful of studies show the importance of non-imprinted genes in parent-of-origin effects (Al Adhami et al., 2015; Mott et al., 2014). For example, QTL analyses of a heterogeneous mouse stock showed that non-imprinted genes mediate parent-of-origin effects on the offspring’s immune system (Mott et al., 2014). Our study describes for the first time a system in which an epigenetic regulator acts in the mother and the father to antagonistically regulate the same non-imprinted genes in the zygote. While the magnitude of effects at many genes may be small, it should be noted that small changes in gene expression can be associated with very different phenotypes (Ruzycki et al., 2015). Our allele-specific mRNA-seq data shows that loss of Pol IV from one parent can impact alleles inherited from both parents in the endosperm. This suggests that Pol IV does not act directly at antagonistic loci but acts instead by regulating other modifiers of gene expression. Yet, this antagonistic regulation can also be viewed through another perspective. Parental conflict can be resolved or paused if both parents can modulate the expression level of a gene or the activity of a pathway to an optimum that is tolerable to each. Pol IV’s role in mediating the antagonistic effects of both parents makes it an ideal system to negotiate optimal gene expression levels. Thus, Pol IV may not be solely an agent of conflict, but also a means to resolving it. Overall, these data suggest that Pol IV is part of a gene regulatory network that is evolving under parental conflict.

## Materials and Methods

### Arabidopsis growth conditions, strains and tissue collection

Plants used in this experiment were grown at 22° C in a Conviron chamber on a 16hr light/8hr dark cycle (120 μM light). The *A. thaliana* mutant used in this study was *nrpd1a-4* (SALK_083051 obtained from ABRC) (Herr et al., 2005) in the Col-0 background. We also utilized *nrpd1a-4* introgressed 4 times into L*er* (Erdmann *et al*, 2017). Endosperm from approximately 100 seeds (7 days after pollination) from at least three siliques was dissected free of embryos and seed coats and pooled for each biological replicate as previously described (Gehring *et al*., 2011). Each biological replicate was collected from crosses that used different individuals as parents. The number of replicates for each experiment was decided based on currently accepted practices in genomic studies. For small RNA experiments, we planned to sample three biological replicates for each genotype. However, we had to discard one of the three pat *nrpd1*+/- sRNA libraries because that library had too few reads.

### mRNA, small RNA and DNA isolation and library construction

Large and small sized RNAs were isolated using the RNAqueous micro RNA isolation kit (Thermo Scientific Fisher). Briefly, endosperm dissected from seeds was collected in lysis buffer and then homogenized with an RNAse-free pellet pestle driven by a Kimble motor. Large and small RNA species were isolated and separated using the manufacturer’s protocol. The RNA concentration of the larger fraction was measured by Qubit. Small RNA libraries were constructed using the NEXTflex sRNA-seq kit V3 (Biooscientific). Final library amplification was carried out for 25 cycles and the libraries were size selected (135-160bp) using a Pippin Prep (Sage Science). mRNA-seq libraries were constructed using a Smart-Seq2 protocol (Picelli et al., 2014). NanoPARE libraries were built as described in Schon *et al* (2018) All libraries were sequenced on the Illumina Hi-Seq 2500.

DNA for bisulfite sequencing was isolated from dissected endosperm at 7 days after pollination using QiaAMP DNA microkit (QIAGEN 56304). Dissected tissue was obtained for two biological replicates for each genotype and incubated overnight in a shaker at 56°C in ATL buffer with Proteinase K. Between 70 and 100ng of endosperm DNA obtained from crosses was subjected to bisulfite treatment using the Methylcode Bisulfite conversion kit (Invitrogen). Analysis of cytosines from chloroplasts with at least ten sequenced reads showed a conversion rate of greater than 98% for all libraries. Bisulfite converted DNA was used to build libraries with the Pico Methyl-Seq library kit (Zymo Research, D5455). 7 cycles of amplification was used for library construction. All libraries were sequenced on the Illumina Hi-Seq 2500 (60bp paired-end).

### Small RNA analysis

Small RNA reads were trimmed with fastq_quality_trimmer (*fastq_quality_trimmer -v -t 20 -l 25*). Cutadapt (Martin, 2011) was used to identify adapter bearing reads of suitable length (*cutadapt -a TGGAATTCTCGGGTGCCAAGG --trimmed-only --quality-base 64 -m 24 -M 40 --max-n 0.5 --too-long-output*). Taking advantage of the random nucleotides on the adapters in NEXTflex kits, we used Prinseq (prinseq-lite-0.20.4) (*prinseq-lite.pl-fastq <infile> -out_format 3-out_good <filename> -derep 1 -log*) to remove PCR duplicates (Schmieder & Edwards, 2011). Filtered reads were aligned to a genome consisting of concatenated Col-0 TAIR10 and L*er* pseudo-genome (Col-0 genome substituted with L*er* SNPs) using Bowtie (v 1.2.2) *bowtie -v 2 --best -p 8 −5 4 −3 4 --sam <index file> <infile.fq>* (Langmead et al., 2009). Reads mapping to L*er* were lifted over to Col-0 using custom scripts (Erdmann et al., 2017). A custom script assign-to-allele was used to identify reads arising from Col-0 or L*er* alleles (https://github.com/clp90/imprinting_analysis/tree/master/helper_scripts). Aligned reads between 21 and 24nt in length were binned based on size. Bedtools was used to count reads in 300-bp windows with 200-bp overlaps and over annotated genes and TEs from Araport 11. DESeq2 (Love et al., 2014) was used to identify features showing differences in small RNA abundance with an adjusted *p-value* of 0.05 or less. One complication with using DESeq2 is that the loss of Pol IV-dependent sRNAs at most loci in *nrpd1*-/- leads to an underestimation of wild-type library size by DESeq2, which increases the proportion of false negatives and undercounts the number of Pol IV-dependent sRNA loci. To allay this effect while analyzing genes, we excluded TEs and applied differential expression analysis to just genic and miRNA loci. These non-TE loci also included Pol IV-independent sRNA loci, which provide an estimate of library size. We separately examined TEs using genic sRNA counts to provide an estimate of library size. ShortStack version 3.8.5 (Axtell, 2013) was also used as an orthogonal approach to identify small RNA peaks from bam alignment file output from Bowtie. Parameters chosen for ShortStack included dicermin= 20, dicermax=25 and a mincov of 0.5 rpm.

### mRNA-seq and NanoPARE analysis

The reads from mRNA-Seq and NanoPARE were trimmed for quality with “*trim_galore -q 25 --phred64 --fastqc --length 20 --stringency 5*” and aligned to the TAIR10 genome using Tophat (v2.1.1) (Kim et al., 2013) using the command *tophat -i 30 -I 3000 --segment-mismatches 1 --segment-length 18 --b2-very-sensitive*. Cuffdiff (v2.1.1) (Trapnell et al., 2013) was used to identify differentially expressed genes for mRNA-Seq data. Aligned NanoPARE read counts at each nucleotide in the genome were counted using Bedtools. Sites with statistical differences in NanoPARE read counts were identified by DESeq2.

### DNA methylation analysis

Reads from Bisulfite sequencing were trimmed for quality using Trim Galore. (https://github.com/FelixKrueger/TrimGalore). Trimmed reads were aligned to the TAIR10 genome using Bismark (Krueger & Andrews, 2011) with parameters set to *-N 1 -L 20 --non_directional*. For this alignment, paired-end reads were treated as single reads. Previously described Bismark methylation extractor and custom scripts (Pignatta et al., 2014, 2015) were used to determine DNA methylation/base and then methylation was calculated for 300 bp windows that overlapped by 200bp. Data from the two biological replicates for each genotype were pooled together for comparison between genotypes. To be included in analysis, windows needed to have at least three overlapping cytosines and a depth of 6 reads/cytosine. Windows that differed between genotypes by 10% CHH, 20% CHG or 30% CG DNA methylation were identified as differentially methylated. Overlapping windows with differential methylation between genotypes were merged into differentially methylated regions. To increase the robustness of our conclusions, we added two data filtering steps. DNA methylation in the endosperm varies between maternal and paternal alleles and bisulfite sequencing is known to potentially enrich for methylated DNA (Ji et al., 2014). Since we were examining the consequences of loss of *NRPD1* in either parent, we could preferentially lose DNA methylation from one set of alleles. This could lead to lower coverage of one set of parental alleles and lead to faulty measurements of DNA methylation. We therefore limited our analyses to genomic regions in which reads arising from the maternally inherited genome accounted for 67%+/- 15% of total DNA reads (based on the fact that that 2/3 of the DNA in endosperm is maternally-inherited). Next, we identified DMRs between the two replicates for each genotype to mark regions where DNA methylation was variable within the same genotype. These regions were excluded from further analysis.

## Data Availability

All high-throughput sequencing data will be available in GEO at GSEXXXX.

## Acknowledgements

We thank the Whitehead Institute Genome Technology Core for high-throughput sequencing services and Dr. Mike Nodine for advice on NanoPARE. This research was supported by National Science Foundation Awards 1453459 and 2101337 to M.G.

## Supplemental Files

**Figure S1:**
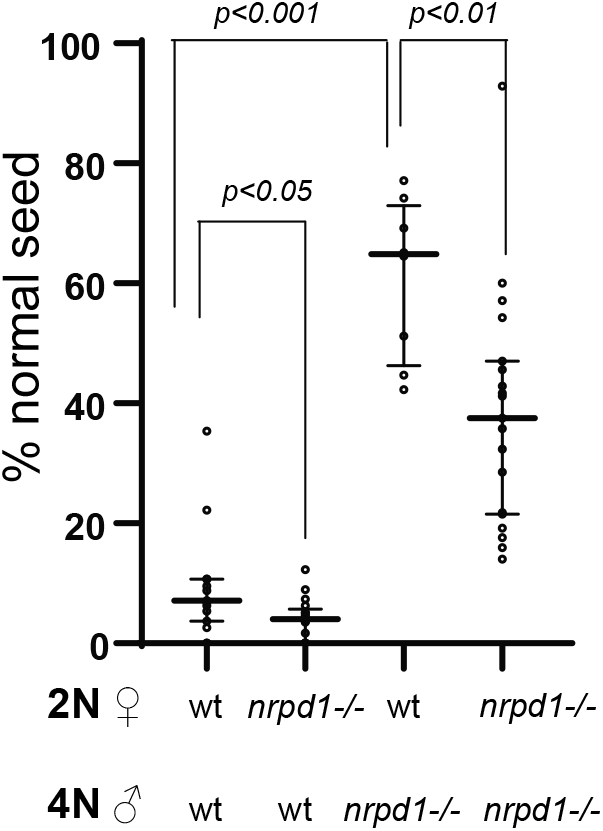
Maternal and paternal Pol IV activity have opposing effects on paternal excess seed abortion. Loss of maternal Pol IV decreases paternal excess seed viability while loss of paternal Pol IV increases seed viability. Each dot in the aligned dot plot represents seed viability from one paternal excess cross (biological replicate). Signficance of difference between indicated crosses was calculated by Wilcox test.

**Figure S2.**
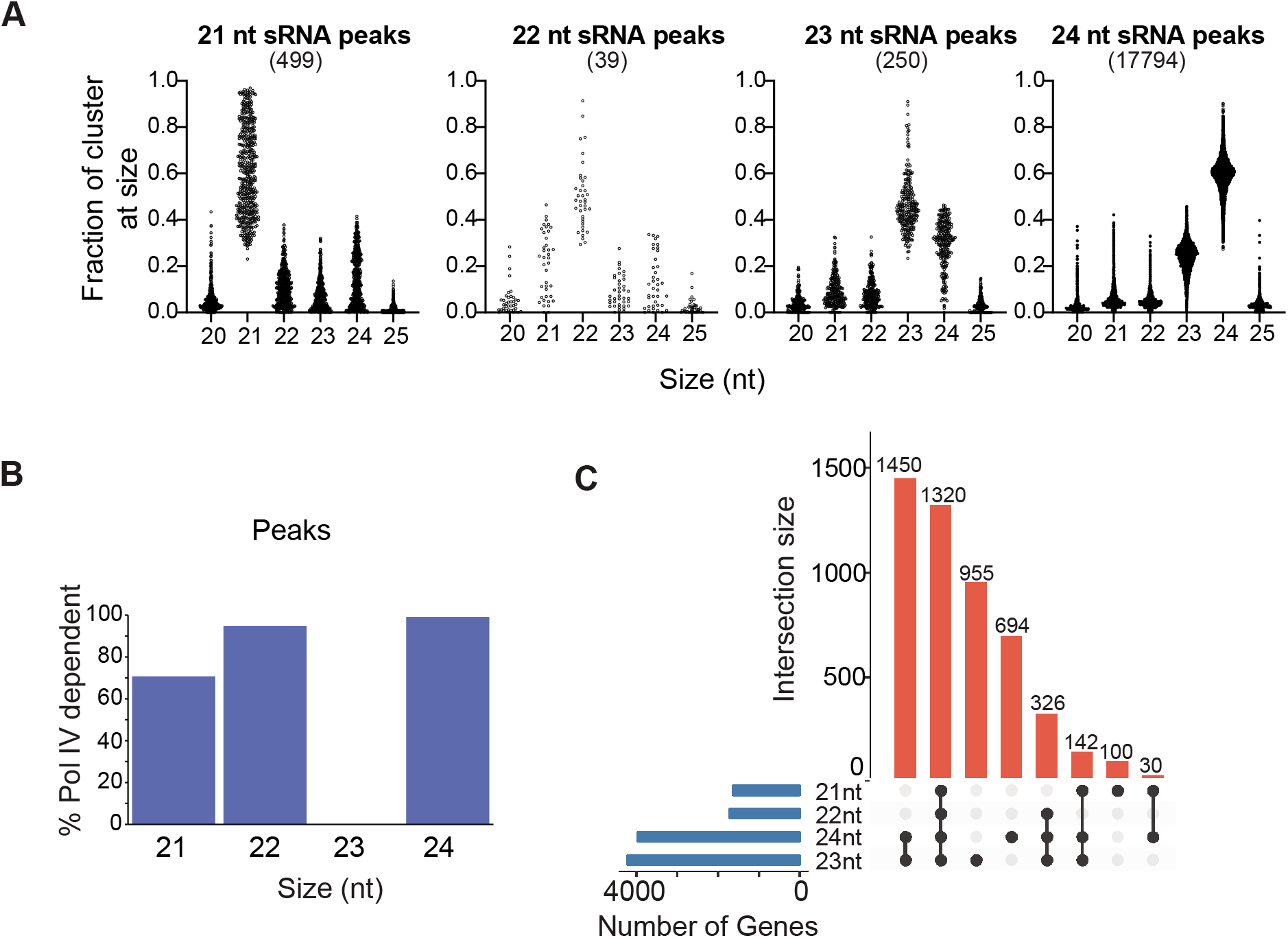
RNA Pol IV is necessary for the production of 21-24 nt sRNAs in the endosperm. **(A)** Size (nt) of all sRNAs in endosperm small RNA peaks dominated by 21, 22, 23, or 24 nt sRNAs. ShortStack was used to call peaks in wild-type (L*er* x Col-0) endosperm. Each peak is grouped into a size class based on the predominant size of the small RNA species in that cluster. Fraction of small RNAs at other sizes at the same peaks are plotted. **(B)** Small RNA peaks of multiple sizes are impacted by loss of *NRPD1*. Genome-wide small RNA coverage over 300 bp windows overlapping by 200 bp was calculated. DESeq2 was used to identify windows with differentially expressed sRNAs between WT and *nrpd1*-/-. Overlapping windows were merged. Peaks overlapping windows with reduced expression in *nrpd1*-/- were classified as Pol IV-dependent peaks. **(C)** Upset plot shows that genes losing sRNAs of one size classes lose sRNAs of other size classes in *nrpd1*-/- endosperm.

**Figure S3:**
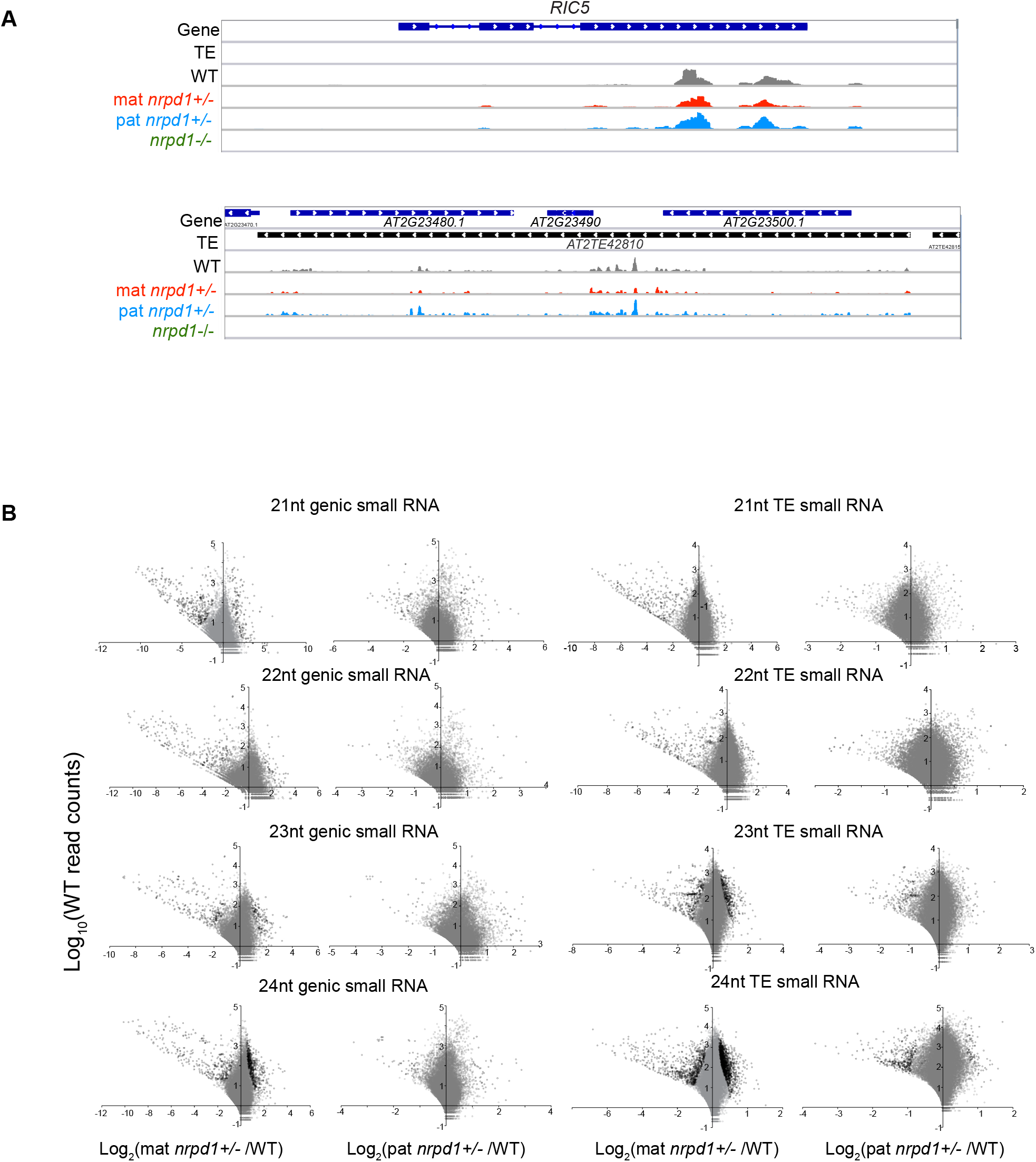
Impact of the loss of maternal and paternal *NRPD1* on the endosperm small RNA population. **(A)** One parent’s copy of *NRPD1* is sufficient for 24 nt sRNA production from genes and TEs at most loci, here exemplifed by *RIC5* and a VANDAL21 copy. **(B)** Examination of 21-24 sRNA over genes and TEs shows that inheriting a maternal mutation in *NRPD1* has a greater impact than inheriting a paternal mutation in *NRPD1*. Loci with differential sRNA expression were identified using DESeq2. Wild-type (WT) read counts represent average read counts per locus across three replicates. Reads mapping to TE insertions were normalized using genic sRNA expression. Black circles represent *padj*≤*0.05*. Gray circles represent *padj*>*0.05*.

**Figure S4:**
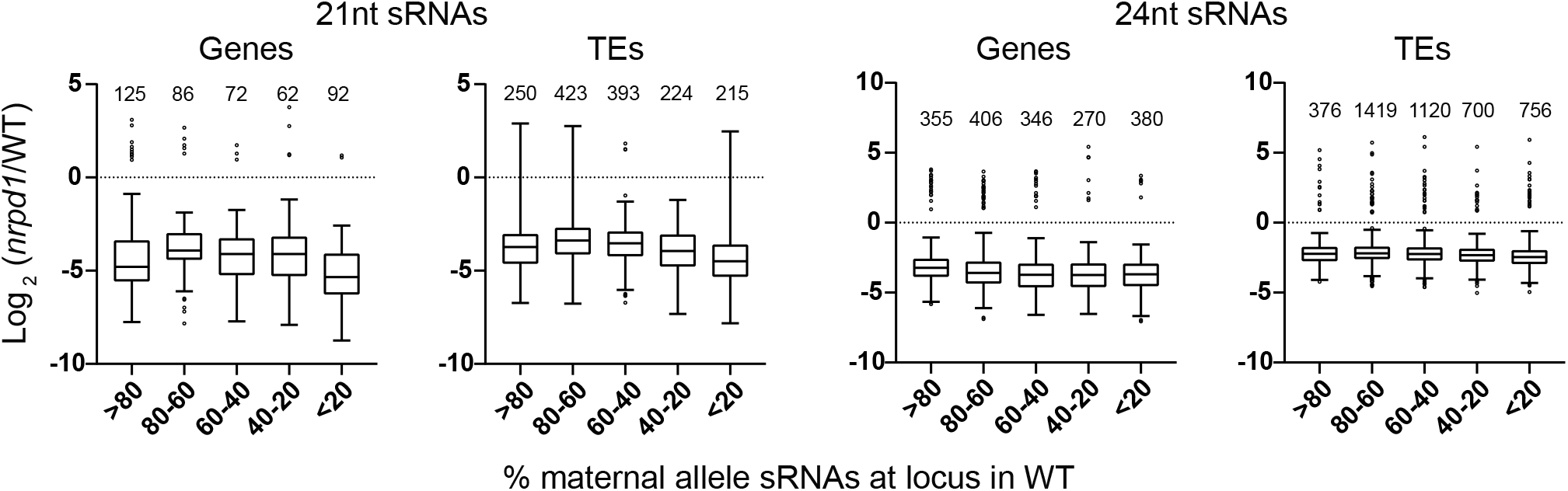
RNA Pol IV-dependent small RNAs arise from both maternal and paternal alleles. SNPs between Col-0 and L*er* were used to identify parental origins of small RNAs arising from genes and transposable elements (TEs). Differentially expressed loci were identified using DESeq2 as described in Figure 1. Loci with a sum of at least ten allele-specific reads in three wild-type L*er* x Col-0 replicates and showing significant differences in 21nt and 24 nt sRNAs in Ler *nrpd1*-/- x Col *nrpdl*-/- endosperm were included. Box plots are Tukey plots. Numbers over box plots are number of loci evaluated.

**Figure S5:**
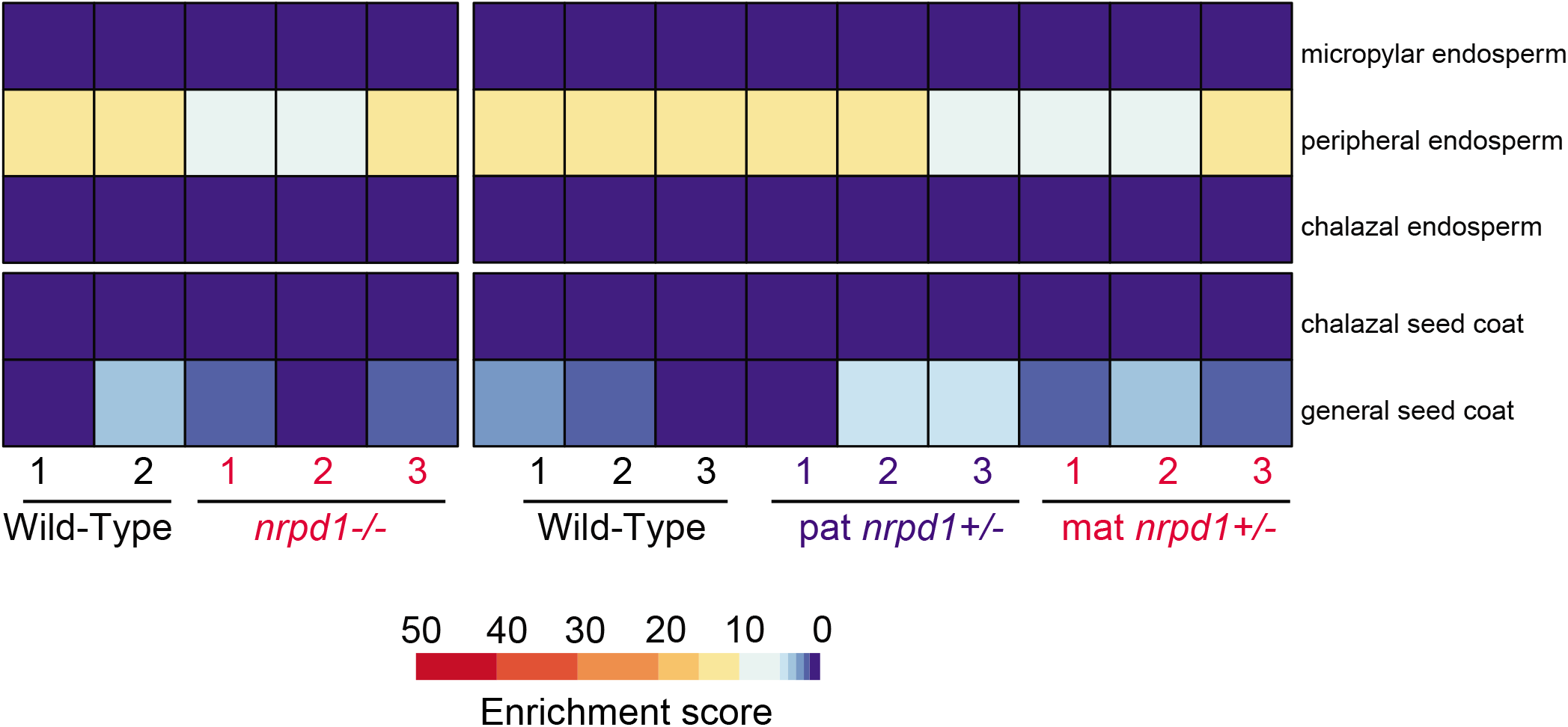
Tissue enrichment in dissected endosperm shows little seed coat contamination. For each RNA-Seq library built with RNA from dissected endosperm, reads overlapping genic loci were counted with Htseq-count. Enrichment of a seed tissue in each sample was then calculated using the Tissue enrichment tool (Schon and Nodine, 2017).

**Figure S6:**
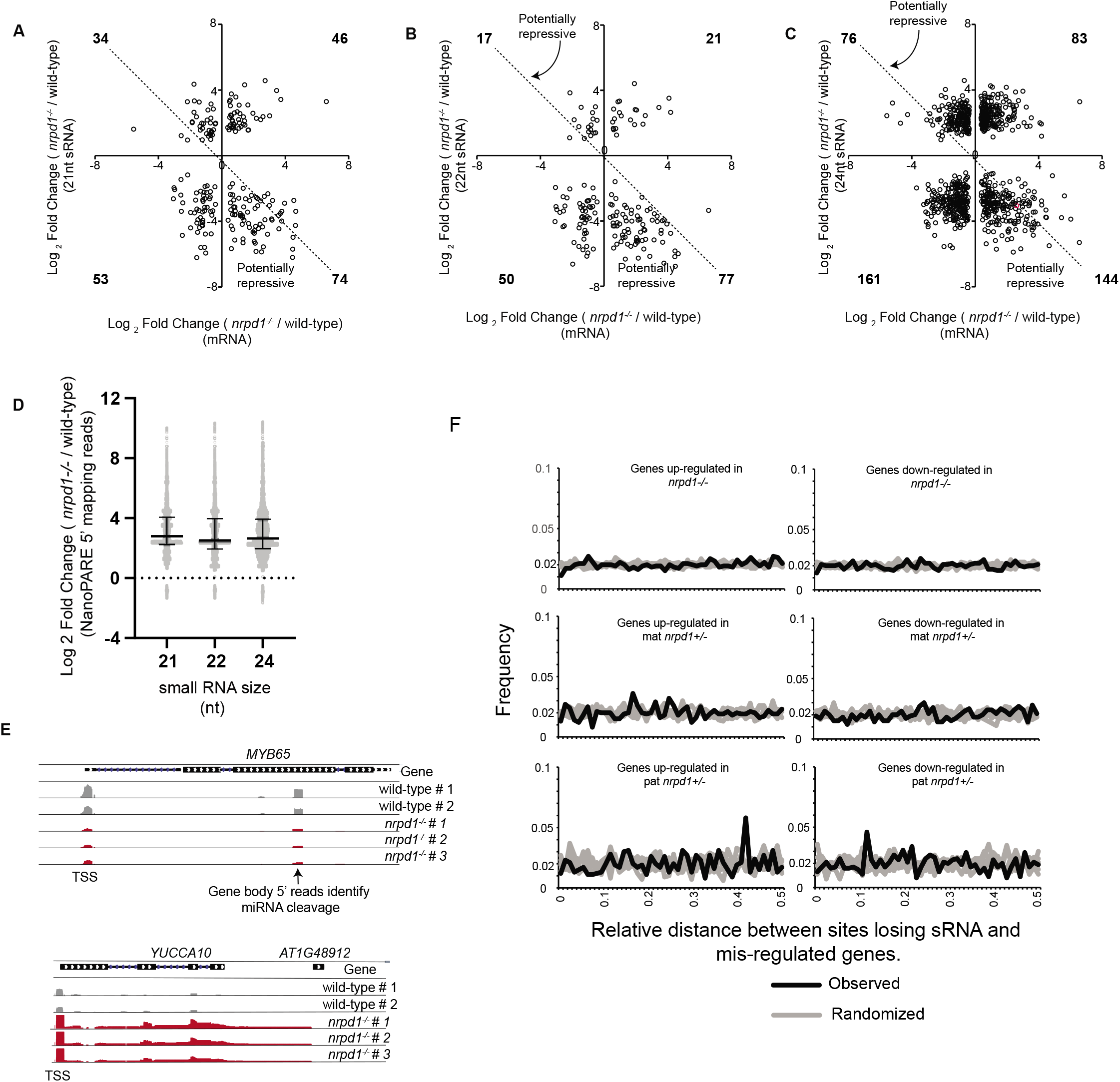
Little relationship between Pol IV sRNAs and gene regulation. **A-C)** Comparisons of genes showing significant differences in 21, 22, 24nt sRNA and mRNA abundance shows that only a subset of genes (lower right quadrant) may be repressed by Pol IV-dependent small RNAs in wild-type. Differences in small RNA abundance between wild-type and *nrpd1*-/- were calculated using DESeq2. Differences in mRNA was calculated using Cuffdiff. Numbers in bold in each quadrant indicate number of genes. **D)** NanoPARE data maps 5’ ends of transcripts and identifies transcriptional start sites (TSS) and cleavage sites within the gene body. Change in mRNA cleavage at genes that show increased mRNA abundance and decreased 21, 22 or 24 nt sRNA size. Coverage of 5’ reads from NanoPARE sequencing was calculated for every nucleotide in the genome. Difference in 5’ read coverage at each nucleotide was calculated for two replicates of wild-type (L*er* x Col-0 endosperm) and three *nrpd1*-/- (L*er nrpd1*-/- x Col-0 *nrpd1*-/-) replicates using DESeq2. Each point plotted on the dot plot represents one nucleotide with differential 5’ reads overlapping a gene. A single gene may thus have more than one 5’ read mapping region. **E)** Examination of NanoPARE data from two replicates of wild-type and *nrpd1*-/- correctly identifies a documented miR159 cleavage site in *MYB65* but identfies no difference in putative cleavage of the *YUCCA10* transcript. *YUCCA10* was chosen as an example because it shows increased mRNA abundance and reduced small RNA abundance in *nrpd1*-/-. **F)**The relative distance metric shows no significant correlation between mis-regulated genes and sites losing small RNAs in nrpd1+/- and nrpd1-/-. Relative distance was calculated using bedtools.Black line indicates relative distance between sites losing sRNA (identified by DESeq2 by examination of readcounts over 300bp windows) and mis-regulated genes. Grey lines represent 5 replicates of equivalent number of random sites in the genome and mis-regulated genes. A uniform frequency of about 0.02 indicates no major correlation between the two data-sets. 5896,1720 and 790 sites lost sRNAs in *nrpd1*-/-, mat *nrpd1*+/- and pat *nrpd1*+/- respectively. The relative choppiness of the the distribution in pat *nrpd1*+/- is likely driven by the smaller number of sites being compared.

**Figure S7:**
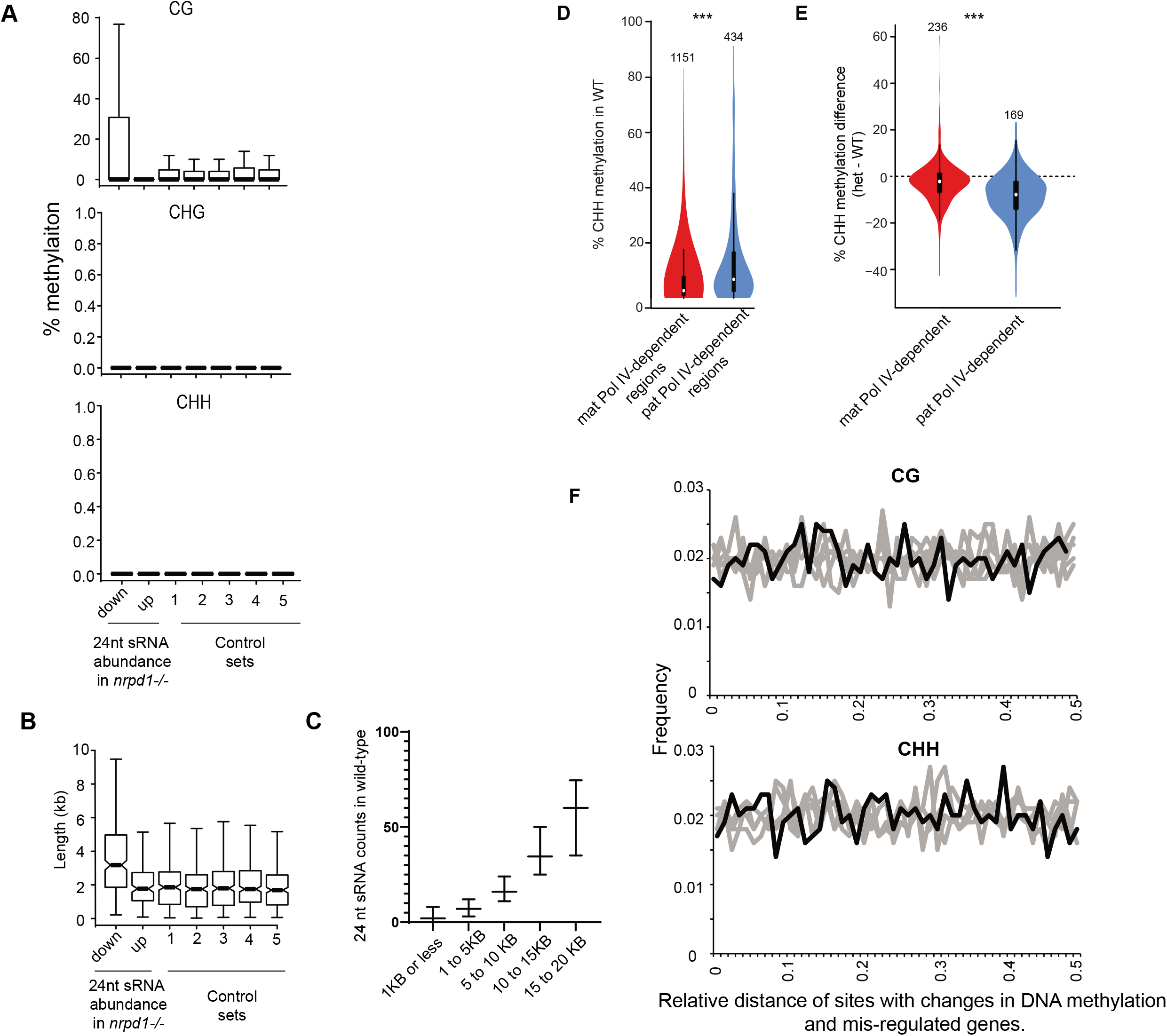
No relationship between DNA methylation changes and genic mis-regulation in *nrpd1*+/-. (**A)** Comparison of methylation in CG, CHG and CHH contexts at individual cytosines within genes in wild-type. Cytosines in the first two columns on the left lie within mis-regulated genes whose sRNA abundances are up or down in *nrpd1*-/-. The control sets include cytosines within five randomly selected sub-set of genes that show no changes in mRNA abundance in *nrpd1*-/-. (**B)** Genes with fewer sRNAs in *nrpd1*-/- and mis-regulated expression tend to be longer. **(C)** Longer genes have more total 24 nt sRNAs in wild-type endosperm. **(D)** Wild-type CHH methylation at sRNA producing sites that are dependent on maternal or paternal Pol IV. Methylation is significantly higher at paternal Pol IV-dependent sites. **(E)** Effects of parental Pol IV loss on CHH methylation at regions with parental Pol IV-dependent sRNAs and greater than 10% CHH methylation in WT. Red, difference between mat *nrpd1*+/- and WT; blue, difference between pat *nrpd1*+/- and WT. Small RNA producing regions impacted in paternal *nrpd1*+/- have higher losses of CHH methylation. For D and E, CHH methylation was calculated for 300bp sliding windows with a 200 bp overlap. CHH methylation windows overlapping windows losing small RNAs in *nrpd1*+/- endosperm were identified and merged using bedtools; maximum CHH methylation among merged windows was used for violin plot. *** represents a statistically significant difference as calculated by Wilcoxon test (p<0.001). Boxplot in the violin plot shows median and inter-quartile range. **(F)** The relative distance metric shows no significant correlation between mis-regulated genes and sites with changes in CG and CHH DNA methylation in mat *nrpd1*+/-. Relative distance was calculated using bedtools.Black line indicates relative distance between mis-regulated genes and sites with differences in DNA methylation between wild-type and mat nrpd1+/- (identified by Bismark). Grey lines represent relative distance betwen 5 replicates of random sites in the genome and mis-regulated genes. A uniform frequency of about 0.02 indicates no major correlation between the two data-sets.

**Figure S8:**
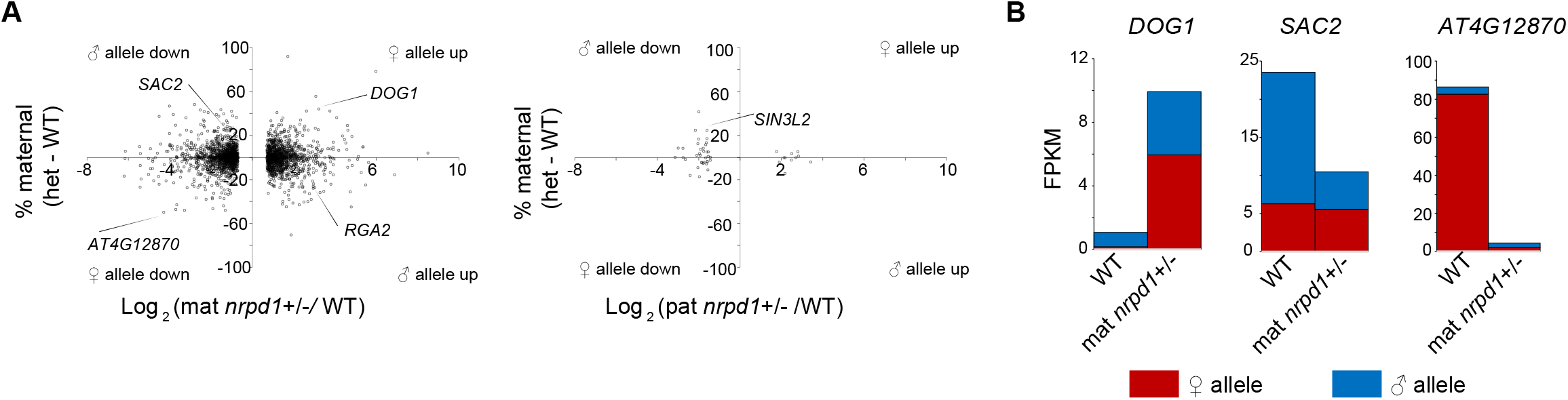
Impact of parental *NRPD1* on maternal and paternal allele contributions to total gene expression. **(A)** Genes were examined to identify those whose expression differences were driven by allele-specific effects. Genes with at least a two-fold, statistically signficant difference in expression between the indicated heterozygote and WT and at least ten allele-specific reads in both genotypes were included. The shift in allelic expression was evaluated by subtracting the % maternal-allele transcripts in WT from the heterozygote. Genes within Col-0 introgressions that remain in L*er nrpdl*-/- plants were excluded from all analyses. **(B)** Examples of genes showing allele-specific impacts upon loss of of maternal Pol IV. FPKM and fold-change in (A) and (B) are from Cuffdiff output.

**Table S1:** List of sequenced libraries.

**Table S2:** DESeq2 output for comparison of 21-24nt sRNA over genes between Wild-type and *nrpd1*-/-, maternal *nrpd1*+/-, paternal *nrpd1*+/-.

**Table S3:** DESeq2 output for comparison of 21-24nt sRNA over transposons between Wild-type and nrpd1-/-, maternal *nrpd1*+/-, paternal *nrpd1*+/-.

**Table S4:** Bedgraph for windows with differences in 21-24nt sRNA between Wild-type and nrpd1-/-, maternal *nrpd1*+/-, paternal *nrpd1*+/-.

**Table S5:** Bed files showing regions differentially methylated between wild-type, mat *nrpd1*+/- and pat *nrpd1*+/- endosperm.

**Table S6:** Cuffdiff output showing genes that are differentially expressed between wild-type, *nrpd1*-/-, mat *nrpd1*+/- and pat *nrpd1*+/-.

**Table S7:** Distance between mis-regulated genes and regions with differences in sRNAs and DNA methylation

